# Effects of Δ9-THC and cannabidiol vapor inhalation in male and female rats

**DOI:** 10.1101/128173

**Authors:** Mehrak Javadi-Paydar, Jacques D. Nguyen, Tony M. Kerr, Yanabel Grant, Sophia A. Vandewater, Maury Cole, Michael A. Taffe

## Abstract

**Rationale:** Previous studies report sex differences in some, but not all, responses to cannabinoids in rats. The majority of studies use parenteral injection, however most human use is via smoke inhalation and, increasingly, vapor inhalation.

**Objectives:** To compare thermoregulatory and locomotor responses to inhaled Δ^9^-tetrahydrocannabinol (THC), cannabidiol (CBD) and their combination using an e-cigarette based model in male and female rats.

**Methods:** Male and female Wistar rats (N=8 per group) were implanted with radiotelemetry devices for the assessment of body temperature and locomotor activity. Animals were then exposed to THC or CBD vapor using a propylene glycol (PG) vehicle. THC dose was adjusted via the concentration in the vehicle (12.5-200 mg/ml) and the CBD (100, 400 mg/mL) dose was also adjusted by varying the inhalation duration (10-40 minutes). Anti-nociception was evaluated using a tail-withdrawal assay following vapor inhalation. Plasma samples obtained following inhalation in different groups of rats were compared for THC content.

**Results:** THC inhalation reduced body temperature and increased tail-withdrawal latency in both sexes equivalently and in a concentration-dependent manner. Female temperature, activity and tail-withdrawal responses to THC did not differ between the estrus and diestrus phases. CBD inhalation alone induced modest hypothermia and suppressed locomotor activity in both males and females. Co-administration of THC with CBD, in a 1:4 ratio, significantly decreased temperature and activity in an approximately additive manner and to similar extent in each sex. Plasma THC varied with the concentration in the PG vehicle but did not differ across rat sex.

**Conclusion:** In summary the inhalation of THC or CBD, alone and in combination, produces approximately equivalent effects in male and female rats. This confirms the efficacy of the e-cigarette based method of THC delivery in female rats.

## Introduction

Human ingestion of *Cannabis sativa* is presumably reinforced by the effects of the phytocannabinoid Δ^9^-tetrahydrocannabinol (THC) on the central nervous system. THC is a partial agonist at the endogenous cannabinoid receptors (CB1 and CB2) and is the major psychoactive component of most recreational cannabis (Burgdorf et al. 2011; ElSohly et al. 2016; Morgan et al. 2010). There may be sex differences in the effects of THC since women reported significantly more dizziness, accompanied by a greater drop in mean arterial pressure (Mathew et al. 2003), and less tachycardia than men after smoking cannabis (Cocchetto et al. 1981). In addition marijuana smoking women exhibit greater deficits in visuospatial memory than their male counterparts during abstinence (Pope et al. 1997). Similarly, THC has been reported to be more potent in female compared with male rodents in producing anti-nociception (Craft et al. 2012; Romero et al. 2002; Tseng and Craft 2001), hypothermia (Borgen et al. 1973; Wiley et al. 2007), and motoric effects (Cohn et al. 1972; Tseng and Craft 2001). On the other hand, no apparent sex differences in subjective intoxication or plasma levels of THC were found in humans after smoking marijuana (Mathew et al. 2003; Miller et al. 1983; Wall et al. 1983) and another study found no sex differences in the effects of THC on impulsivity in human subjects (McDonald et al. 2003). In laboratory findings, male rodents may be more sensitive than females to the hyperphagic effect of cannabinoid agonists (Diaz et al. 2009) and females may be more sensitive than males to the adverse effects of escalating adolescent THC exposure on emotional behavior and stress reactivity in adulthood (Rubino et al. 2008). The fact that cannabinoids bind with greater affinity to CB1 receptors in female compared with male rats (Craft et al. 2012), and that CB1 expression differs between normally cycling females and either male or ovarectomized female rats (Riebe et al. 2010) suggests a sex-specific mechanism and further indicates that observed sex differences may vary across the estrous cycle. Thus it remains of significant interest to determine similarities and differences in the effects of THC between sexes.

Humans typically smoke cannabis, and more recently are turning to noncombusted inhalation techniques (Jones et al. 2016; Morean et al. 2015; Varlet et al. 2016), yet almost all preclinical studies of sex differences with cannabinoids have involved systemic injection such as intraperitoneal (Wiley et al. 2007) or intravenous (Martin et al. 1991) administration. The route of administration of cannabinoids, and particularly THC, can cause variability in the pharmacokinetic, pharmacodynamic and behavioral effects in humans and non-human animals (Fried 1976; Fried and Nieman 1973; Manwell et al. 2014; Naef et al. 2004; Niyuhire et al. 2007; Wilson et al. 2002; Wilson et al. 2006). Parenteral administration of cannabinoid agonists suppresses spontaneous activity, decreases nociception, induces hypothermia, and increases catalepsy in rodents of both sexes (Compton et al. 1993; Martin et al. 1991; Wiley et al. 2007) thus these measures predominate in rodent models of the effects of THC. Inhalation delivery of THC with a custom metered-dose inhaler confirmed comparable degrees of antinociception, hypothermia, and catalepsy in mice exposed to the THC aerosol (Lichtman et al. 2000, Wilson, 2002#57) compared with parenteral injection of THC. In addition, THC levels in the blood and brains of mice were similar following either marijuana smoke inhalation exposure or intravenous injection of THC, further confirming that inhalation delivery of THC is a viable method to elicit cannabinoid-typical pharmacological effects (Lichtman et al. 2001).

An additional phytocannabinoid of recent interest (see (Boggs et al. 2018) for review), cannabidiol (CBD), does not activate the CB1 and CB2 receptors (Pertwee 2006; 2008) but has shown activity in opposition to the effects of CB1/CB2 receptor agonists in some findings (Morgan et al. 2010; Radwan et al. 2009; Wright et al. 2013). On the other hand, CBD does not oppose, and may enhance, hypothermia caused by THC, i.p. when administered in 1:1-1:3 THC:CBD ratios in rats (Taffe et al. 2015). Additionally, CBD-enhanced discriminative stimulus effects of THC have been reported in monkeys, albeit at very low THC:CBD ratios (McMahon 2016). It remains unknown how CBD may alter the effects of THC when administered by inhalation in laboratory species.

Our recent study (Nguyen et al. 2016b) validated a new method of rat cannabinoid inhalation to model current human use of e-cigarettes to administer cannabinoids. That prior study included a limited sex comparison that appeared to demonstrate greater female hypothermia following THC inhalation. Thus a new study was designed to more fully compare the effects of THC inhalation between male and female rats on nociception, thermoregulation, spontaneous activity and plasma THC levels. Experiments were included to assess THC effects during estrus and diestrus to determine any possible contributions of the estrous cycle. The effects of inhaled CBD on thermoregulation and activity were determined in isolation and in combination with THC.

## Materials and methods

### Subjects

Age-matched groups of Wistar rats (Charles River, NC, USA) were housed in humidity and temperature-controlled (23±1 °C) vivaria on 12:12 hour light:dark cycles. Animals entered the laboratory at ∼10 weeks of age. Animals had *ad libitum* access to food and water in their home cages. All procedures were conducted in the animals’ dark (active) cycle under protocols approved by the Institutional Care and Use Committees of The Scripps Research Institute and in a manner consistent with the *Guide for the Care and Use of Laboratory Animals (National Research Council (U.S.). Committee for the Update of the Guide for the Care and Use of Laboratory Animals. et al. 2011).*

### Radiotelemetry

Male (N=8) and female (N=8) Wistar rats were anesthetized with an isoflurane/oxygen vapor mixture (isoflurane 5% induction, 1-3% maintenance) and sterile radiotelemetry transmitters (Data Sciences International, St. Paul, MN; TA-F40) were implanted in the abdominal cavity through an incision along the abdominal midline posterior to the xyphoid space. Absorbable sutures were used to close the abdominal muscle incision and the skin incision was closed with the tissue adhesive. For the first three days of the recovery period, an antibiotic (cefazolin; 0.4 g/ml; 2.0 ml/kg, s.c.) and an analgesic (flunixin; 2.5 mg/ml; 2.0 ml/kg, s.c.) were administered daily. A minimum of 7 days were allowed for surgical recovery prior to starting any experiments. This minimally invasive technique is useful for characterizing increases and decreases in both body temperature and locomotor activity consequent to drug exposure in both monkeys (Crean et al. 2007; Taffe 2011; 2012; Taffe et al. 2006) and rats (Miller et al. 2013a; Miller et al. 2013b).

Activity and temperature responses were evaluated in clean standard plastic home cages (thin layer of bedding) in a dark testing room, separate from the vivarium, during the (vivarium) dark cycle. In some experiments (detailed below) animals were habituated in recording chambers, moved to the vapor inhalation chamber for the exposure and then returned to the separate recording chamber. In other experiments, telemetry plates under the vapor inhalation chambers were used throughout the recording. Radiotelemetry transmissions were collected via telemetry receiver plates placed under the cages as previously described (Aarde et al. 2013; Miller et al. 2013b; Wright et al. 2012).

### Inhalation Apparatus

Sealed exposure chambers were modified from the 259mm X 234mm X 209mm Allentown, Inc (Allentown, NJ) rat cage to regulate airflow and the delivery of vaporized drug to rats using e-cigarette cartridges (Protank 3 Atomizer by Kanger Tech; Shenzhen Kanger Technology Co.,LTD; Fuyong Town, Shenzhen, China) as has been previously described (Nguyen et al. 2016a; Nguyen et al. 2016b). An e-vape controller (Model SSV-1; La Jolla Alcohol Research, Inc, La Jolla, CA, USA) was triggered to deliver the scheduled series of puffs by a computerized controller designed by the equipment manufacturer (Control Cube 1; La Jolla Alcohol Research, Inc, La Jolla, CA, USA). The chamber air was vacuum controlled by a chamber exhaust valve (i.e., a “pull” system) to flow room ambient air through an intake valve at ∼1 L per minute. This also functioned to ensure that vapor entered the chamber on each device triggering event. The vapor stream was integrated with the ambient air stream once triggered.

### Nociception Assay

Tail withdrawal anti-nociception was assessed using a water bath (Bransonic(®) CPXH Ultrasonic Baths, Danbury, CT) maintained at 48, 50 or 52 °C. Three different temperature conditions were evaluated in the event of any non-linearities in the effect of THC across rat sex. The latency to withdraw the tail was measured using a stopwatch and a cutoff of 15 seconds was used to avoid any possible tissue damage (Wakley and Craft 2011; Wakley et al. 2014a). Tail withdrawal was assessed 60 minutes after the initiation of inhalation (i.e. 30 minutes after termination of each session).

### Drugs

The inhalation exposure was to Δ^9^-tetrahydrocannabinol (THC; 12.5, 25, 50, 100, 200 mg/mL) or cannabidiol (100, 400 mg/mL) in propylene glycol (PG) vehicle. Four 10-s vapor puffs were delivered with 2-s intervals every 5 minutes, which resulted in use of approximately 0.125 ml in a 40 minutes exposure session (Nguyen et al. 2016b). The vacuum was turned off for the 4 minute, 12 second interval between vapor deliveries and then turned up to ∼3-5 L/minutes at the conclusion of sessions for ∼5 minutes to facilitate complete chamber clearance for subject removal. THC was suspended in a vehicle of 95% ethanol, Cremophor EL and saline in a 1:1:8 ratio, for intraperitoneal injection. The THC was provided by the U.S. National Institute on Drug Abuse Drug Supply Program and cannabidiol was obtained from Cayman Chemical (Ann Arbor, MI).

### Determination of estrous stage

The stage of the estrous cycle was determined using interpretation of vaginal cytology. Unstained vaginal smears were viewed immediately upon collection via pipet lavage. Proestrus was indicated when cells were predominantly nucleated epithelial cells. A predominance of cornified epithelial cells classified the estrus stage. Metestrus was recognized by scattered, nucleated or cornified epithelial cells and leukocytes, and diestrus was classified by classic leukocytes in combination with various larger round epithelial cells. (Freeman 1988; Goldman et al. 2007). Vaginal samples were taken in the evening prior to the day of experiment to facilitate planning the dosing and a second smear was obtained ∼ 1 hour before THC inhalation to confirm. The estrus and diestrus stages were selected for this experiment because a previous study reported the largest stage-related differential effect of 5 mg/kg THC in estrus versus diestrus in gonadally intact, cycling females (Craft and Leitl 2008).

### Plasma THC/CBD analysis

For single time point studies blood samples were collected (∼500-1000 ul) via jugular needle insertion under anesthesia with an isoflurane/oxygen vapor mixture (isoflurane 5% induction, 1–3% maintenance). For serial blood sampling (200 ul; 35, 60, 120 and 240 minutes after vapor initiation) rats were prepared with chronic intravenous catheters as previously described (Aarde et al. 2015; Aarde et al. 2017). Catheter function was assessed prior to starting a series of experiments and only animals with a functional catheter were used; a two week recovery interval was imposed prior to any further experimentation for multiple-sampling procedures. Plasma THC content was quantified using fast liquid chromatography/mass spectrometry (LC/MS) adapted from (Irimia et al. 2015; Lacroix and Saussereau 2012; Nguyen et al. 2017). Fifty uL of plasma were mixed with 50 uL of deuterated internal standard (100ng/ml CBD-d3 and THC-d3; Cerilliant), and cannabinoids were extracted into 300 uL acetonitrile and 600 uL of chloroform and then dried. Samples were reconstituted in 100 uL of an acetonitrile/methanol/water (2:1:1) mixture. Separation was performed on an Agilent LC1100 using an Eclipse XDB-C18 column (3.5um, 2.1mm x 100mm) using gradient elution with water and methanol, both with 0.2 % formic acid (300 uL/min; 73-90%). Cannabinoids were quantified using an Agilent MSD6180 single quadrupole using electrospray ionization and selected ion monitoring [CBD (m/z=313.7), CBD-d3 (m/z=316.7), THC (m/z=313.7) and THC-d3 (m/z=316.7)]. Calibration curves were conducted for each assay at a concentration range of 0-200 ng/mL and observed correlation coefficients were 0.999.

### Experiments

A minimum 7 day interval separated all active THC dosing for a given individual throughout the following experimental conditions.

#### Experiment 1

The first experiment was conducted in experimentally naïve animals to determine the effect of altering THC concentration in the PG. Animals were placed in individual telemetry recording cages for at least 30 minutes of habituation prior to the start of inhalation. This initial 30-minute telemetry data were used as the pre-treatment baseline for statistical analysis purposes. Then, rats were transferred in pairs to separate inhalation chambers for vapor exposure and thereafter returned to their recording cages (this approach was used for Experiments 2-4). In this first experiment, telemetry recording was conducted following dosing conditions of PG or THC (12.5, 25, 50, 100 mg/mL) with the order randomized in pairs. Recording continued for up to 4 hours after the start of vapor inhalation. Subsequent experiments were conducted in the same groups of male and female animals with the test conditions randomized within experiment as described below.

#### Experiment 2

After Experiment 1, the female telemetry group was recorded during the estrus and diestrus phases of the estrous cycle. First the female rats were recorded following 30 minutes of inhalation of THC 50 mg/mL (versus PG) in estrus and diestrus phases for four total treatment conditions. They were next assessed following 40 minutes of inhalation of THC 25 mg/mL versus PG to further explore the potentially more rapid recovery in estrus identified after the 50 mg/mL THC inhalation.

#### Experiment 3

After Experiment 1, the male group was evaluated for responses to 30 min of inhalation of PG, CBD (100 mg/ml), THC (200 mg/mL) or the combination in randomized order.

#### Experiment 4

Next, both male and female groups were evaluated for responses to CBD inhalation. In this case, the dosing conditions were CBD (100 mg/mL) for three different inhalation durations (10, 20, 40 minutes), or PG for a 20 minute duration, in a randomized order.

#### Experiment 5

After Experiment 4, telemetry recordings were taken for male and female rat groups following exposure to PG, CBD (100 mg/mL), THC (25 mg/mL) or the combination for 30 minutes. For this study, the telemetry recording was conducted throughout vapor inhalation, thus animals were dosed singly and all recordings were from rats in the inhalation chamber. All 8 animals within the male/female groups were run simultaneously (different inhalation conditions) on a given day. Male and female groups were evaluated on different days.

#### Experiment 6

Nociception was assessed in both male and female groups 60 minutes after the initiation of inhalation of PG or THC (100 mg/mL) for 30 minutes in randomized order. Following this study, the female group was evaluated for nociception after inhalation of PG or THC (100 mg/mL) for 30 minutes in the estrus and diestrus phases of the estrous cycle, in randomized order.

#### Experiment 7

Finally, telemetry recordings for both male and female rats were obtained following inhalation of PG, CBD (400 mg/ml), THC (100 mg/mL) or the combination for 30 minutes in randomized dosing order. This represented the same 1:4 THC:CBD ratio evaluated in Experiment 5 but at a higher overall dose. Animals were dosed and recorded as described in Experiment 5.

#### Experiment 8

Pharmacokinetic experiments were conducted in additional groups of male and female Wistar rats. The first group of male (N=8) and female (N=8) Wistar rats were 20 weeks of age at the start of this study. These rats arrived in the lab as a cohort and were previously implanted with jugular catheters, for methods see (Aarde et al. 2015; Aarde et al. 2017); one male and one female rat did not survive catheter surgery. Intact rats with patent catheters at 20 weeks of age (2 out of 7 male and 3 out of 7 female rats) were used for serial plasma collection at 35, 60, 120 and 240 minutes after the start of inhalation of THC Vapor (100 mg/ml, 30 min). Then, the same total group of rats (7 per sex; male mean weight 424.1 g SEM 17.0, female mean weight 256.2 g SEM 5.1) were exposed to 30 min of inhalation of THC in three concentrations (25, 100, 200 mg/mL) with single jugular blood withdrawals (∼0.5 mL) obtained post-session (i.e., 35 minutes after the start of inhalation) and at 60 minutes after the start of inhalation for 6 total sessions. THC exposure was conducted no more frequently than weekly for single blood withdrawals and biweekly for multiple blood draws (Diehl et al. 2001). A second group of male and female Wistar rats, 38 weeks of age at the start of this study, were used for determining plasma levels after i.p. injection of THC. These animals arrived in the lab as a cohort (N=8 per sex) and were previously exposed to doses of oxycodone, heroin, methadone and buprenorphine via vapor inhalation and parenteral injection over an interval of 14 weeks for anti-nociception experiments (one male rat died after an oxycodone challenge thus N=7 for this study). The PK study was conducted starting 12 weeks after the end of the prior studies at 38 weeks of age (male mean weight 706.3 g SEM 22.8; female mean weight 337.9 g SEM 13.6). The rats were divided into two independent groups (3-4 rats per sex) and were injected with one of two THC doses (3, 10 mg/kg, i.p.) followed by a single blood collection obtained 30 minutes post-injection.

### Data Analysis

The body temperature and activity rate (counts per minute) were collected via the radiotelemetry system on a 5-minute schedule and analyzed in 30 minute averages (in the graphs the time point refers to the ending time, i.e. 60 = average of 35-60 minute samples). Any missing temperature values were interpolated from preceding and succeeding values. Telemetry data were analyzed with Analysis of Variance (ANOVA) with repeated measures factors for the Drug Inhalation Condition, the Time post-initiation of vapor and estrus phase where relevant. Tail-withdrawal latencies were analyzed by ANOVA with repeated measures factors of Drug Treatment Condition and Water Bath temperature and between-groups factor of sex. Plasma levels of THC were analyzed with between-groups ANOVA due to the design with factors for Time post-initiation of inhalation, drug Concentration in the PG and/or sex, as described in the Results for specific experiments. Any significant effects within group were followed with post-hoc analysis using Tukey correction for all multi-level, and Sidak correction for any two-level, comparisons. All analysis used Prism 7 for Windows (v. 7.03; GraphPad Software, Inc, San Diego CA).

## Results

### Experiment 1: Effect of Vapor Inhalation of THC (12.5-100 mg/mL) in Male and Female Rats

#### Females

The body *temperature* of female rats was decreased by THC inhalation in a concentration dependent manner (**Figure 1**). The ANOVA confirmed significant effects of Time Post-initiation [F (7, 49) = 18.49; P < 0.0001], of Vapor Conditions [F (4, 28) = 13.44; P < 0.0001] and of the interaction of factors [F (28, 196) = 9.10; P < 0.0001]. Temperature was significantly different from the baseline after inhalation of 25 mg/mL (120-240 minutes), 50 mg/mL (90-210 minutes) or 100 mg/mL (60-240 minutes) THC but not after the 12.5 mg/mL or PG inhalation. Temperature did not differ compared with PG in the 12.5 or 25 mg/mL concentrations, but the post-hoc test confirmed significant differences from vehicle after 50 mg/mL (60-150, 210 minutes) or 100 mg/mL (60-150 minutes) THC inhalation. Temperature was also significantly different 60-90 minutes after the start of inhalation of 50 mg/mL versus 100 mg/mL. In addition, the temperature was significantly lower compared with the 12.5 mg/mL condition following 50 mg/mL (60-90 minutes) or 100 mg/mL (60-120 minutes) and lower than the 25 mg/mL following 50 mg/mL (60-120 minutes) or 100 mg/mL (60-120 minutes).

**Figure 1:**
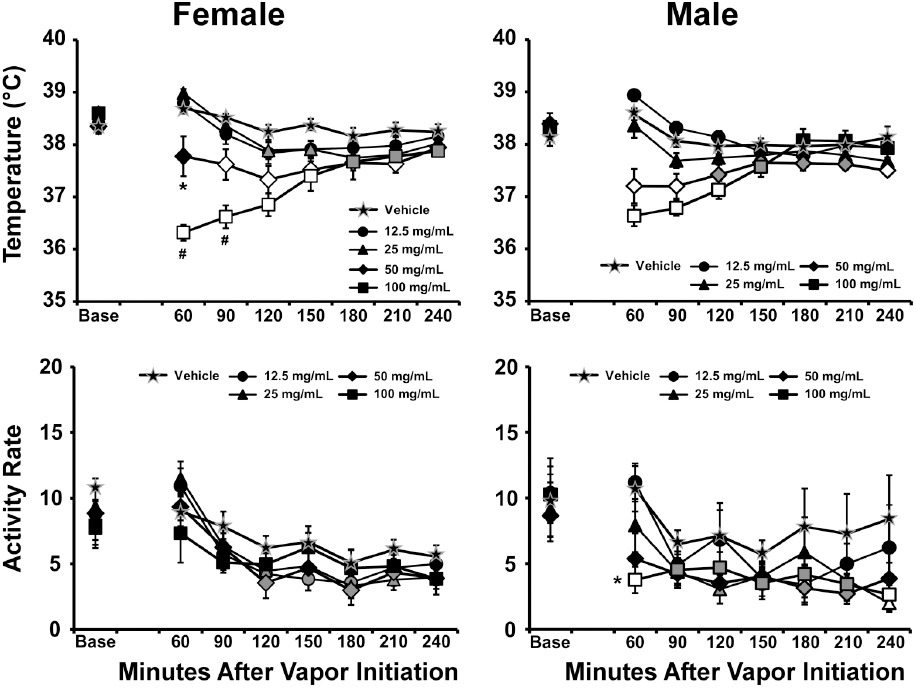
Mean female (N=8; ±SEM) and male (N=8; ±SEM) body temperatures and activity following inhalation exposure to the polyethylene glycol vehicle (PG) or THC (12.5-100 mg/mL in PG) vapor for 30 minutes. Open symbols indicate a significant difference from both vehicle at a given time-point and the within-treatment baseline, while shaded symbols indicate a significant difference from the baseline only. A significant difference from the vehicle condition (only) at a given time is indicated by * and from the 50 mg/mL condition by #. Base=baseline value.

The *activity* of female rats was significantly affected by Time Post-initiation [F (7, 49) = 26.78; P < 0.0001] but not by Vapor Condition or by the interaction of factors. The marginal mean post-hoc analysis of the Time factor confirmed significantly lower activity rates compared either with the baseline (90-240 minutes post-initiation) or with the 60 minute time-point (90-240 minutes).

### Males

The body *temperature* of male rats was also decreased by THC inhalation (**Figure 1**) and the ANOVA confirmed significant effects of Time Post-initiation [F (7, 49) = 8.65; P < 0.0001], of the five Vapor Conditions [F (4, 28) = 11.09; P < 0.0001] and of the interaction of factors [F (28, 196) = 6.45; P < 0.0001]. The temperature was significantly different from the baseline after 50 mg/mL (60-240 minutes) or 100 mg/mL (60-150 minutes) THC inhalation but not after the 12.5 mg/mL, 25 mg/mL or PG conditions. The post-hoc test confirmed significant differences from vehicle after 50 mg/mL (60-90, 240 minutes) or 100 mg/mL (60-120 minutes) THC inhalation. Temperature did not differ compared with PG in the 12.5 or 25 mg/mL concentrations at any time post-initiation of vapor. In addition, the temperature of male rats was significantly lower compared with the 12.5 mg/mL condition following 50 mg/mL (60-120 minutes) or 100 mg/mL (60-120 minutes) and lower than the 25 mg/mL following 50 mg/mL (60 minutes) or 100 mg/mL (60-120 minutes).

The *activity* of male rats was significantly affected by Time Post-initiation [F (7, 49) = 16.53; P < 0.0001] and by Vapor Condition [F (4, 28) = 3.48; P <0.05] but not by the interaction of factors. The marginal mean analysis confirmed that significantly less activity was observed after 50 mg/mL or 100 mg/mL THC inhalation compared with the vehicle. Likewise the marginal mean post-hoc analysis of the Time factor confirmed significantly lower activity rates were observed compared with the baseline (90-240 minutes post-initiation) and compared with the 60 minute time-point (90-240 minutes).

Follow-up analysis compared the impact of vapor inhalation in the 60 min time point compared with the baseline across groups to directly compare the sexes. This three-way ANOVA first confirmed that temperature was significantly affected by inhalation dose condition [F (4, 4) = 31.95; P <0.0001], by timepoint [F (1, 4) = 21.15; P <0.0001] and by the interaction of dose with time [F (4, 4) = 36.09; P<0.0001] but not by sex or the interaction of sex with any other factor. A similar analysis failed to confirm any main or interacting effects of sex, dosing condition or time on activity rate.

### Experiment 2: Impact of Estrous Phase on the Effects of Inhaled THC

#### THC (50 mg/mL; 30 minutes)

Mean body temperature was significantly lowered by THC inhalation (**Figure 2**) as confirmed by a significant effect of Drug / estrous condition [F (3, 21) = 19.15; P < 0.0001], Time Post-Initiation [F (9, 63) = 13.12; P < 0.0001], and the interaction of factors [F (27, 189) = 3.42; P < 0.0001]. Body temperature was significantly different from the baseline value after 30 minutes exposure to 50 mg/ml THC (60-210 minutes) for estrus phase and (60-270 minutes) for diestrus phase. The post-hoc test also confirmed that temperature differed from the corresponding vehicle after 30 minutes exposure to 50 mg/ml THC (60-180 minutes) for estrus phase and (60-300 minutes) for diestrus phase. Finally temperature was significantly different between estrus and diestrus phase, after 240 minutes post-exposure to THC 50 mg/ml.

**Figure 2:**
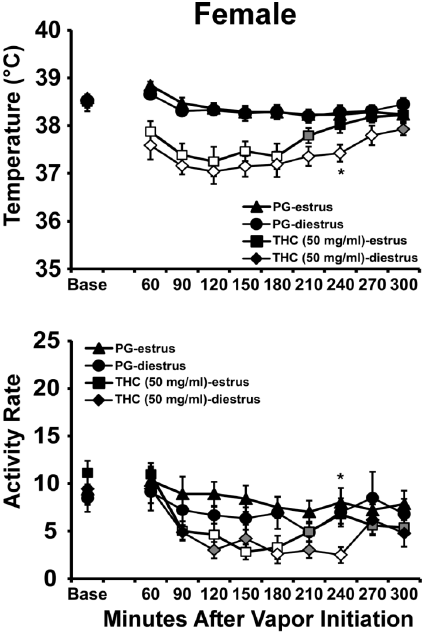
Mean (N=8; ±SEM) temperature and activity following vapor inhalation of the PG or THC (50 mg/mL in PG) for 30 minutes in estrus and diestrus stages. Open symbols indicate a significant difference from both vehicle and the baseline, while shaded symbols indicate a significant difference from the baseline only; A significant difference between estrus and diestrus phase in THC 50 mg/ml at a given time is indicated by *

The ANOVA confirmed significant effects of Time Post-initiation [F (9, 63) = 13.80, P<0.0001] and of Vapor Condition [F (3, 21) = 5.00; P<0.01] on activity rate. The Tukey post-hoc test confirmed that activity was significantly different from the baseline after 30 minutes exposure to 50 mg/ml THC (90-210 and 270-300 minutes) for estrus phase and (120-240 minutes) for diestrus phase. The post-hoc test also confirmed that activity differed from the corresponding vehicle after 30 minutes exposure to 50 mg/ml THC (120-180 minutes) for estrus phase and (180, 240 minutes) for diestrus phase. Finally activity was significantly different between estrus and diestrus phase, after 240 minutes post-exposure to THC 50 mg/ml.***THC (25 mg/mL; 40 minutes)***: In this study N=6 completed the estrus evaluations and N=7 completed the diestrus evaluations due to scheduling constraints (**Figure 3**). The female rats’ body temperature was again significantly affected by THC dose condition [F (3, 22) = 5.66; P < 0.01], by Time Post-Initiation [F (9, 198) = 14.94; P < 0.0001] and by the interaction of factors [F (27, 198) = 2.83; P < 0.0001]. The post-hoc test further confirmed that temperature was significantly different from the baseline value after 40 minutes of inhalation 25 mg/ml THC (90-180 minutes) for estrus phase and (90-150 minutes) for diestrus phase. The post-hoc test also confirmed that temperature differed from the corresponding vehicle after 40 minutes of exposure to 25 mg/ml THC (90-180 minutes) for both estrus and diestrus phases. Finally temperature was not significantly different between estrus and diestrus phase within either drug condition. Exposure to THC 12.5 mg/ml for 40 minutes did not affect body temperature and activity in either estrus or diestrus phases.

**Figure 3:**
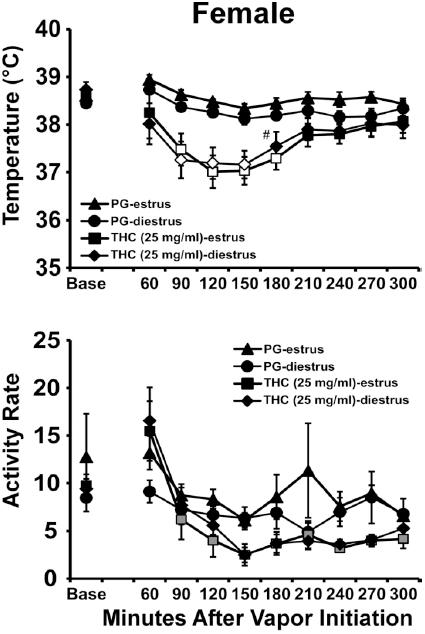
Mean (N=6-7; ±SEM) temperature and activity following vapor inhalation of the PG or THC (25 mg/mL in PG) for 40 minutes in estrus and diestrus stages. Open symbols indicate a significant difference from both vehicle and the baseline, while shaded symbols indicate a significant difference from the baseline only; A significant difference from the vehicle condition at a given time and phase is indicated by #.

Activity rate was significantly affected by Time Post-Initiation [F (9, 198) = 14.52; P < 0.0001]. Locomotor activity was significantly different from the baseline value after 40 minutes exposure to 25 mg/ml THC (90-300 minutes) for diestrus phase. Activity rate was not significantly different between estrus and diestrus phase, after exposure to THC 25 mg/ml.

### Experiment 3: Vapor Inhalation of THC with Cannabidiol (CBD)

#### Males

Males were next evaluated in conditions of PG versus CBD (100 mg/mL) for 30 minutes in randomized order and then PG, THC (200 mg/mL) and THC (200 mg/mL) + CBD (100 mg/mL) for 30 minutes in randomized order. Preliminary analysis identified no differences between the first and second PG conditions and thus the second one was used for analysis purposes. Body temperature was again decreased by cannabinoid inhalation (**Figure 4**) and the ANOVA confirmed significant effects of Time Post-initiation [F (7, 49) = 40.83; P<0.0001], of the four Vapor Conditions [F (3, 21) = 37.3; P<0.0001] and of the interaction of factors [F (21, 147) = 11.88; P<0.0001]. The Tukey post-hoc test confirmed that body temperature was significantly different from the baseline after PG (150-240 minutes), CBD 100 mg/mL (60-240 minutes), THC 200 mg/mL (60-240 minutes), and after THC 200 mg/mL + CBD 100 mg/mL (60-240 minutes) inhalation. Correspondingly, body temperature was significantly lower than the PG inhalation condition following CBD 100 mg/mL (60-120 minutes), THC 200 mg/mL (60-150 minutes), and after THC 200 mg/mL + CBD 100 mg/mL (60-180 minutes) inhalation. In addition, significant differences from the CBD 100 mg/mL condition were confirmed for THC 200 mg/mL (60-150, 240 minutes), and after THC 200 mg/mL + CBD 100 mg/mL (60-150, 240 minutes) inhalation. Body temperature was never different between the THC 200 mg/mL and THC 200 mg/mL + CBD 100 mg/mL inhalation conditions.

**Figure 4:**
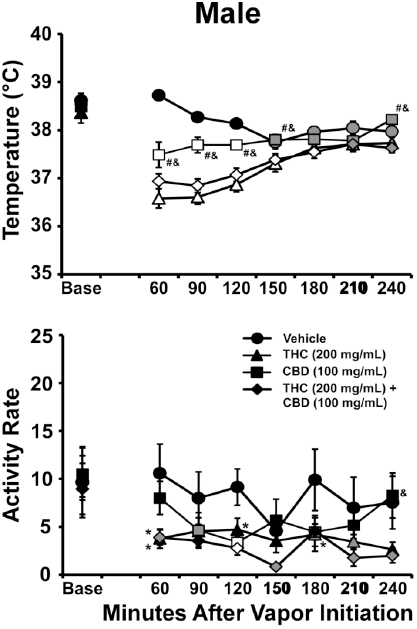
Mean (N=8; ±SEM) temperature and activity following vapor inhalation of the PG, THC (200 mg/mL in PG), cannabidiol (CBD; 100 mg/mL) or the THC /CBD combination. Open symbols indicate a significant difference from both vehicle at a given time-point and the within-treatment baseline, while shaded symbols indicate a significant difference from the baseline only. A significant difference from vehicle condition (only) is indicated by *, from the THC (200 mg/mL) condition at a given time by # and from the THC (200 mg/mL)+CBD (100 mg/mL) condition by &. Base=baseline value.

Activity rate was also altered by drug inhalation conditions and the ANOVA confirmed significant effects of Time Post-initiation [F (7, 49) = 5.47; P=0.0001], of the four Vapor Conditions [F (3, 21) = 6.06; P<0.005] and of the interaction of factors [F (21, 147) = 2.12; P<0.01]. The Tukey post-hoc test confirmed that activity was significantly different from the baseline after CBD 100 mg/mL (90-120,180 minutes), THC 200 mg/mL (210 minutes), and after THC 200 mg/mL + CBD 100 mg/mL (120-150, 210-240 minutes) inhalation. Activity was significantly lower than the PG inhalation condition following CBD 100 mg/mL (120 minutes), THC 200 mg/mL (60, 120, 180 minutes), and after THC 200 mg/mL + CBD 100 mg/mL (60, 120 minutes) inhalation. A significant difference from the CBD 100 mg/mL condition was confirmed for the THC 200 mg/mL + CBD 100 mg/mL condition 240 minutes after the start of inhalation. Activity was never significantly different between the THC 200 mg/mL and THC 200 mg/mL + CBD 100 mg/mL inhalation conditions.

### Experiment 4: Cannabidiol Duration/Response

#### Females

The body *temperature* of female rats was decreased by CBD (**Figure 5**) and the ANOVA confirmed significant effects of Time Post-initiation [F (7, 49) = 5.84, P<0.0001] and of the interaction of Time Post-initiation with Vapor Condition [F (21, 147) = 3.05; P<0.0001]. The Tukey post-hoc test confirmed that temperature was significantly lower compared with the baseline following inhalation of CBD for 20 minutes (120-180 minutes after the start of inhalation) and for 40 minutes (60-180 minutes). Temperature was also significantly lower compared with the PG condition following inhalation of CBD for 40 minutes (60-90 minutes after the start of inhalation).

**Figure 5:**
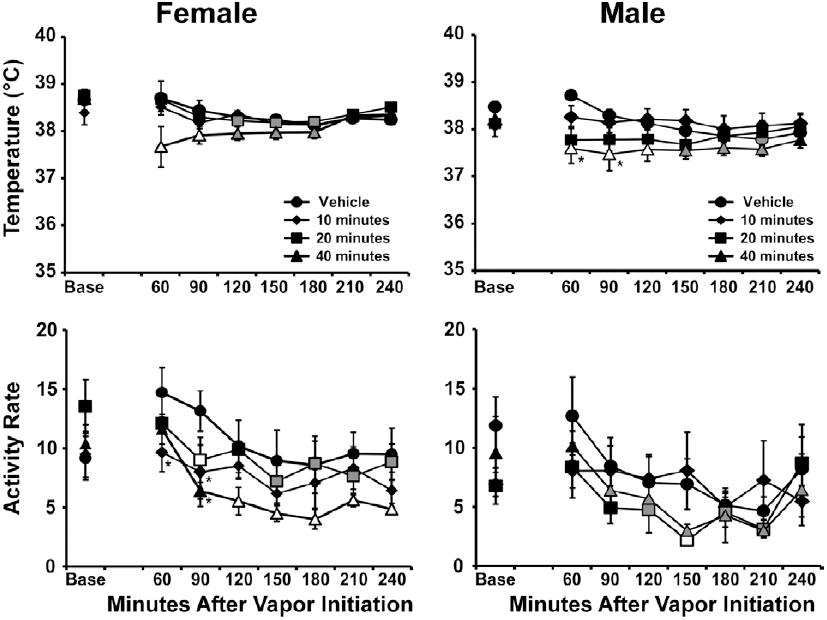
Mean female (N=8; ±SEM) and male (N=8; ±SEM) body temperatures and activity following inhalation exposure to the polyethylene glycol vehicle (PG) or CBD (100 mg/mL in PG) vapor for 10, 20 and 40 minutes vapor inhalation. Open symbols indicate a significant difference from both vehicle and the baseline, while shaded symbols indicate a significant difference from the baseline only. A significant difference from the vehicle condition (only) at a given time is indicated by *.

The ANOVA also confirmed significant effects of Time Post-initiation [F (7, 49) = 19.39, P<0.0001] and of the interaction of Time Post-initiation with Vapor Condition [F (21, 147) = 1.90; P<0.05] on *activity* rate. The Tukey post-hoc test confirmed that activity was significantly lower compared with the baseline following inhalation of CBD for 20 minutes (90, 150-240 minutes after the start of inhalation) or 40 minutes (120-240 minutes after the start of inhalation). Activity rate was also significantly lower compared with the PG condition following inhalation of CBD for 10 minutes (60-90 minutes after the start of inhalation) as well as CBD for 20 minutes (90 minutes); and lower compared with the baseline following inhalation of CBD for 40 minutes (90-240 minutes).

#### Males

The body *temperature* of male rats (**Figure 5**) was again altered by CBD and the ANOVA confirmed significant effects of Time Post-initiation [F (7, 49) = 3.52; P<0.005] and of the interaction of Time Post-initiation with Vapor Condition [F (21, 147) = 2.24; P<0.005]. The Tukey post-hoc test confirmed that temperature was significantly lower compared with the baseline following inhalation of PG (180-210 minutes after the start of inhalation) as well as CBD for 40 minutes (60-210 minutes).

Temperature was also significantly lower compared with the PG condition following inhalation of CBD for 20 minutes (60-90 minutes after the start of inhalation) or 40 minutes (60-120 minutes) and lower compared with the CBD 10 minutes condition following inhalation of CBD for 20 minutes (60, 150 minutes after the start of inhalation) or 40 minutes (60-150, 210 minutes).

Male rat *activity* rate was also affected by vapor inhalation and the ANOVA confirmed significant effects of Time Post-initiation [F (7, 49) = 11.59; P<0.0001] and of Vapor Condition [F (3, 21) = 4.63; P<0.05], but not of the interaction of factors, on activity rate. The post-hoc of the marginal means confirmed that activity was lower in the 20 minutes CBD inhalation condition compared with PG and across treatment conditions, activity was lower than the baseline (120-210 minutes after the start of inhalation) or the 40 minutes (90-240 minutes) time-points.

### Experiment 5: Effects of Threshold THC (25 mg/mL) + CBD (100 mg/mL) Combination

#### Females

The body temperature of female rats was decreased by drug inhalation (**Figure 6**) and the ANOVA confirmed significant effects of Time Post-initiation [F (8, 56) = 22.53; P<0.0001], of Vapor Condition [F (3, 21) = 5.19; P<0.01] and of the interaction of factors [F (24, 168) = 4.23; P<0.0001]. The Tukey post-hoc test confirmed that body temperature was significantly different from the baseline after PG (180-240 minutes), CBD 100 mg/mL (30-240 minutes), THC 25 mg/mL (90-240 minutes), and after THC 25 mg/mL + CBD 100 mg/mL (30-240 minutes) inhalation. Correspondingly, body temperature was significantly lower than the PG inhalation condition following CBD 100 mg/mL (30-60 minutes), THC 25 mg/mL (60, 120-210 minutes), and after THC 25 mg/mL + CBD 100 mg/mL (30-240 minutes) inhalation. In addition, significant differences from the CBD 100 mg/mL condition were confirmed for THC 25 mg/mL (60, 210 minutes), and for THC 25 mg/mL + CBD 100 mg/mL (60, 120-240 minutes) inhalation. Body temperature was also different between the THC 25 mg/mL and THC 25 mg/mL + CBD 100 mg/mL, (60 minutes) after inhalation conditions.

**Figure 6:**
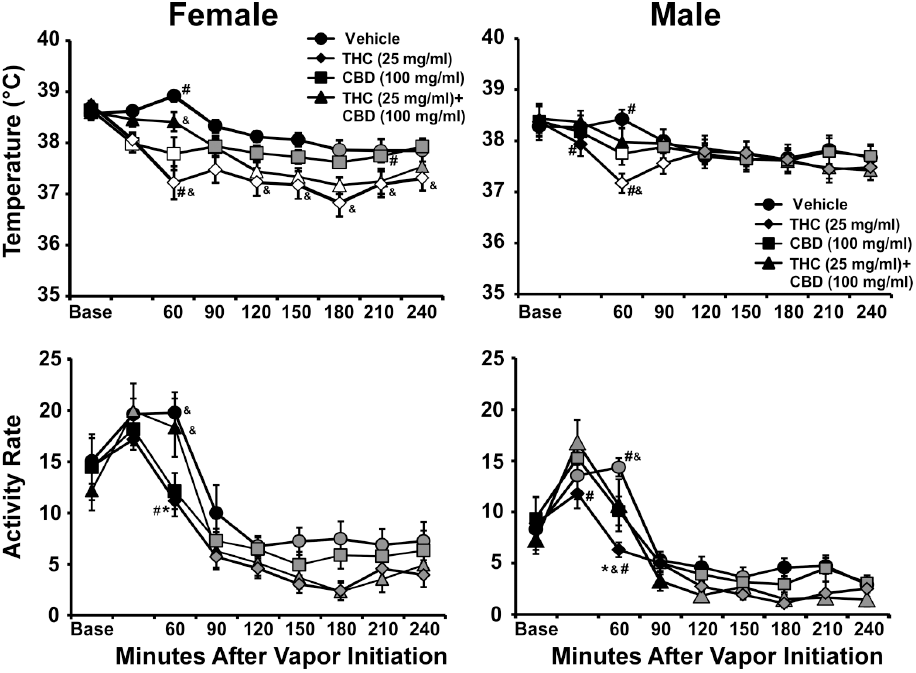
Mean female (N=8; ±SEM) and male (N=8; ±SEM) temperature and activity following vapor inhalation of the PG, THC (25 mg/mL in PG), cannabidiol (CBD; 100 mg/mL) or the THC/CBD combination. Open symbols indicate a significant difference from both vehicle and the baseline, while shaded symbols indicate a significant difference from the baseline only. A significant difference from the vehicle condition (only) at a given time is indicated by *, a difference from THC (25 mg/ml) by # and a significant difference from CBD (100 mg/ml) by &.

The ANOVA confirmed significant effects of Time Post-initiation [F (8, 56) = 70.27; P=0.0001] on activity rate. The Tukey post-hoc test confirmed that activity was significantly different from the baseline after PG (120-240 minutes), CBD 100 mg/mL (90-240 minutes), THC 25 mg/mL (30, 120-240 minutes), and after THC 25 mg/mL + CBD 100 mg/mL (90-240 minutes) inhalation. Activity was significantly lower than the PG inhalation condition following CBD 100 mg/mL (60 minutes) and after THC 25 mg/mL + CBD 100 mg/mL (60 minutes) inhalation. A significant difference from the CBD 100 mg/mL condition was confirmed for THC 25 mg/mL 60 minutes after the start of inhalation and activity was also significantly different between the THC 25 mg/mL and THC 25 mg/mL + CBD 100 mg/mL conditions, 60 minutes after the start of inhalation.

#### Males

The ANOVA confirmed significant effects of Time Post-initiation [F (8, 56) = 15.16; P<0.0001], and of the interaction of factors [F (24, 168) = 3.36; P<0.0001] on body temperature (**Figure 6**). The Tukey post-hoc test confirmed that body temperature was significantly different from the baseline after PG (120-180, 240 minutes), CBD 100 mg/mL (60-240 minutes), THC 25 mg/mL (90-240 minutes), and after THC 25 mg/mL + CBD 100 mg/mL (60-240 minutes) inhalation. Correspondingly, body temperature was significantly lower than the PG inhalation condition following CBD 100 mg/mL (60 minutes), THC 25 mg/mL (60 minutes), and after THC 25 mg/mL + CBD 100 mg/mL (60-90 minutes) inhalation. In addition, significant differences from the CBD 100 mg/mL condition were confirmed for THC 25 mg/mL + CBD 100 mg/mL (60 minutes) inhalation. Body temperature was also different between the THC 25 mg/mL and THC 25 mg/mL + CBD 100 mg/mL (30-60 minutes) conditions.

The ANOVA confirmed significant effects of Time Post-initiation [F (8, 56) = 31.10; P=0.0001], of Vapor Condition [F (3, 21) = 4.97; P<0.01] and of the interaction of factors [F (24, 168) = 2.02; P<0.01] on activity rate. The Tukey post-hoc test confirmed that activity was significantly different from the baseline after PG (30-60, 150, 240 minutes), CBD 100 mg/mL (30, 90-240 minutes), THC 25 mg/mL (30, 120-240 minutes), and after THC 25 mg/mL + CBD 100 mg/mL (120-240 minutes) inhalation. Activity was significantly lower than the PG inhalation condition following CBD 100 mg/mL (60 minutes), THC 25 mg/mL (60 minutes), and after THC 25 mg/mL + CBD 100 mg/mL (60 minutes) inhalation. There was a significant difference from the CBD 100 mg/mL condition which was confirmed 60 minutes after the start of inhalation in the THC 25 mg/mL + CBD 100 mg/mL condition. Activity also differed significantly between the THC 25 mg/mL and THC 25 mg/mL + CBD 100 mg/mL (30-60 minutes) conditions.

### Experiment 7: Effects of High Dose THC (100 mg/mL) + CBD (400 mg/mL) Combination

The initial THC+CBD combination in male rats was conducted using a higher THC concentration (200 mg/mL) and a 2:1 THC:CBD ratio and the second study used a 1:4 THC:CBD ratio at lower concentrations. To further explore potentially additive effects, male (N=8) and female rats (N=8) completed a study of the effects of inhalation of PG, CBD (400 mg/mL), THC (100 mg/mL) versus THC (100 mg/mL) + CBD (400 mg/mL) for 30 minutes in randomized order.

#### Females

The body temperature of female rats was decreased by drug inhalation (**Figure 7**) and the ANOVA confirmed significant effects of Time Post-initiation [F (8, 56) = 24.01; P<0.0001], of Vapor Condition [F (3, 21) = 15.3; P<0.0001] and of the interaction of factors [F (24, 168) = 5.19; P<0.0001]. The Tukey post-hoc test confirmed that body temperature was significantly different from the baseline after PG (120-240 minutes), CBD 400 mg/mL (30-240 minutes), THC 100 mg/mL (30-240 minutes), and after THC 100 mg/mL + CBD 400 mg/mL (30-240 minutes) inhalation. Correspondingly, body temperature was significantly lower than the PG inhalation condition following CBD 400 mg/mL (30-180, 240 minutes), THC 100 mg/mL (30-210 minutes), and after THC 100 mg/mL + CBD 400 mg/mL (30-240 minutes) inhalation. In addition, significant differences from the CBD 400 mg/mL condition were confirmed for THC 100 mg/mL (90-150 minutes), and for THC 100 mg/mL + CBD 400 mg/mL (60-240 minutes) inhalation. Body temperature was also different between the THC 100 mg/mL and THC 100 mg/mL + CBD 400 mg/mL, (30-60, 240 minutes) after inhalation conditions.

**Figure 7:**
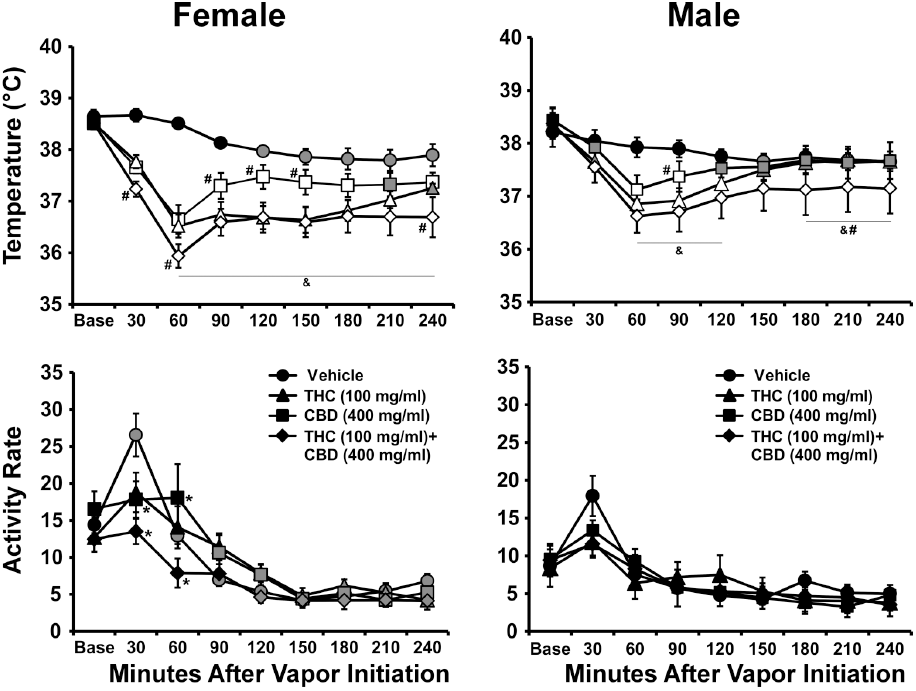
Mean female (N=8; ±SEM) and male (N=8; ±SEM) temperature and activity following vapor inhalation of the PG, THC (100 mg/mL in PG), cannabidiol (CBD; 400 mg/mL) or the THC/CBD combination. Open symbols indicate a significant difference from both vehicle and the baseline while shaded symbols indicate a significant difference from the baseline only. A significant difference from the vehicle condition (only) at a given time is indicated by *, a difference from THC (100 mg/ml) by # and a significant difference from CBD (400 mg/ml) by &.

The ANOVA confirmed significant effects of Time Post-initiation [F (8, 56) = 34.82; P<0.0001] and the interaction of Time with Dosing condition [F (24, 168) = 3.26; P<0.0001] on activity rate. The Tukey post-hoc test confirmed that activity was significantly different from the baseline after PG (30, 90-240 minutes), CBD 400 mg/mL (90-240 minutes), THC 100 mg/mL (30, 150-240 minutes), and after THC 100 mg/mL + CBD 400 mg/mL (120-240 minutes) inhalation. Activity was also significantly different than the PG inhalation condition following CBD 400 mg/mL (30-60 minutes) and after THC 100 mg/mL + CBD 400 mg/mL (30-60 minutes) inhalation. A significant difference from the CBD 400 mg/mL condition was confirmed for THC 100 mg/mL+ CBD 400 mg/mL 60 minutes after the start of inhalation. Activity was also significantly different between the THC 100 mg/mL and THC 100 mg/mL + CBD 400 mg/mL, 30-60 minutes after initiation of inhalation.

#### Males

The ANOVA confirmed significant effects of Time Post-initiation [F (8, 56) = 12.25; P<0.0001], and of the interaction of factors [F (24, 168) = 3.64; P<0.0001] on body temperature (**Figure 7**). The Tukey post-hoc test confirmed that body temperature was significantly different from the baseline after PG (150, 210-240 minutes), CBD 400 mg/mL (30-240 minutes), THC 100 mg/mL (30-240 minutes), and after THC 100 mg/mL + CBD 400 mg/mL (30-240 minutes) inhalation. Correspondingly, body temperature was significantly lower than the PG inhalation condition following CBD 400 mg/mL (60-90 minutes), THC 100 mg/mL (60-120 minutes), and after THC 100 mg/mL + CBD 400 mg/mL (30-240 minutes) inhalation. In addition, significant differences from the CBD 400 mg/mL condition were confirmed for THC 100 mg/mL (90 minutes) and THC 100 mg/mL + CBD 400 mg/mL (60-120, 180-240 minutes) inhalation. Body temperature was also different between the THC 100 mg/mL and THC 100 mg/mL + CBD 400 mg/mL, (180-240 minutes) after inhalation conditions.

The ANOVA confirmed significant effects of Time Post-initiation [F (8, 56) = 13.86; P<0.0001], but not of Vapor Condition or of the interaction of factors, on activity rate. The marginal mean post-hoc test confirmed that across vapor conditions the activity rate was significantly higher than all other time points at 30 min after the start of vapor and significantly lower than baseline activity 150-240 minutes after the start of inhalation.

### No Time-Dependent Difference in the Effect of THC

To assess possible plasticity in the response to the inhalation of THC across dosing conditions, a further analysis compared the temperature response to 100 mg/mL THC (for 30 minutes) in the first experiment and the high-dose combination study **(Figure 8)**. Analysis found no difference in the initial male and female body temperature responses across time points. The female temperature at Time 2 returned to baseline more slowly with significant differences from each other group/time 150-180 min post-initiation and from the males at Time 1 at 150-210 minutes post-initiation. The 2-way ANOVA confirmed that body temperature was significantly affected by sex/dose condition [F (3, 28) = 3.51; P < 0.05], by Time Post-Initiation [F (6, 168) = 35.10; P < 0.0001] and by the interaction of factors [F (18, 168) = 2.39; P < 0.01]. The Tukey post-hoc test further confirmed that temperature was significantly different from males at Time 1 at 150-210 minutes and from males at Time 2 at 150-180 minutes. The mean body weight of the females was 61% of that of the males at Time 1 and 52% at Time 2. Female weight increased by 43% from Time 1 to Time 2 and the males increased by 66%.

**Figure 8:**
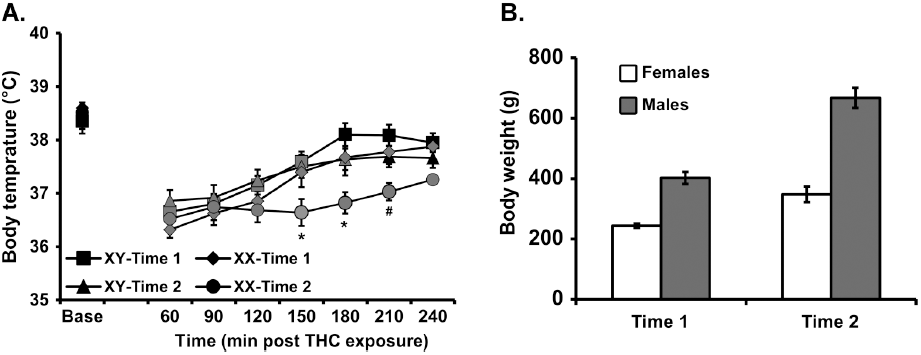
A) Mean male and female (N=8 per group; ±SEM) temperature following vapor inhalation of THC (100 mg/mL in PG) for 30 minutes in Experiments 1 (Time 1) and 7 (Time 2). Shaded symbols indicate a significant difference from the baseline. A significant difference from all other groups/Times is indicated by * and from the males at Time 1 by #. B) Mean body weight for male and female groups at Time 1 and Time 2.

### Experiment 6: Anti-nociceptive Effect of THC

THC inhalation decreased sensitivity to a noxious stimulus in both males and female rats and this effect was similar across different water bath temperature condition (**Figure 9**). The initial 3-way ANOVA confirmed that tail withdrawal latency was significantly affected by sex [F (1, 2) = 16.3; P < 0.0001], THC/PG condition [F (1, 2) = 59.8; P < 0.0001] and water bath temperature [F (2, 2) = 140; P < 0.0001]. The analysis also confirmed a significant interaction of THC treatment condition and sex [F (1, 2) = 5.01; P < 0.05] and an interaction of water temperature with THC/PG treatment [F (2, 2) = 3.48; P<0.05]. The Tukey post-hoc test including all possible comparisons confirmed that latency was significantly longer after THC inhalation for female rats when evaluated at 48°C and 50°C. Significantly longer latencies were confirmed for male versus female rat tail withdrawal from 48°C water after PG inhalation. Follow-up two-way ANOVA were conducted within sex groups to further parse these effects. This analysis first confirmed significant effects of Water Temperature and Vapor condition, but not the interaction, for female [F (2, 14) = 106.9; P<0.0001, F (1, 7) = 65.27; P<0.0001] and male [F (2, 14) =168.6; P<0.0001, F (1, 7) = 12.87; P<0.01] rats. The marginal mean post-hoc analysis further confirmed significant differences between PG and THC inhalation, and between all three water bath temperatures for each sex. Since the inhalation conditions were randomized for the full tail-withdrawal study, not all estrus stages were captured for all female rats. Thus, additional sessions were conducted to complete estrus (PG N=4; THC N=5) and diestrus (PG N=6; THC N=5) evaluations in the 50 °C water-bath only. In this case the tail withdrawal latency was significantly slowed by THC inhalation, but not estrous stage, as confirmed by a significant effect THC/PG inhalation condition [F (1, 7) = 51.51; P < 0.001] without effect of estrous stage or the interaction of factors. The post-hoc test also confirmed an anti-nociceptive effect of THC in each of the estrus and diestrus phases.

**Figure 9:**
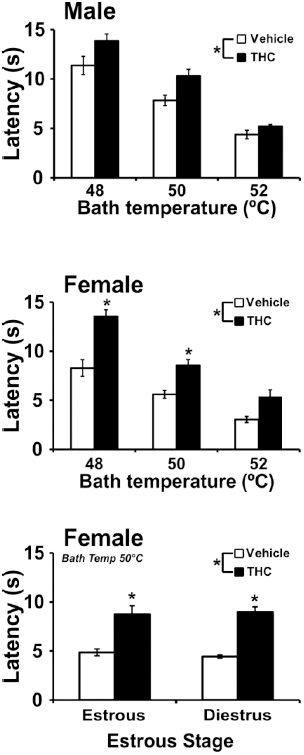
Mean (N=8; ±SEM) tail withdrawal latency following vapor inhalation of the PG or THC (100 mg/mL in PG) for 30 minutes. A significant difference between the vehicle and THC conditions is indicated by *.

### Experiment 8: Pharmacokinetics

Analysis of the single time-point blood draws that were obtained after 30 minutes of inhalation of three THC concentrations (25, 100, 200 mg/mL) showed that plasma THC varied by concentration in the vehicle but not by sex (**Figure 10A**). The statistical analyses confirmed a significant effect of concentration on plasma THC for the 35 minute [F (2, 30) = 23.18; P<0.0001] and 60 minute samples [F (2, 34) = 19.79; P<0.0001], but there was no significant effect of sex or of the interaction of factors. The post-hoc tests also confirmed that for each time point significantly higher plasma THC was observed after inhalation of 200 mg/mL compared with either lower concentration in each sex. Analysis of the plasma sampled serially after a single inhalation session (30 minutes; THC 100 mg/mL) in the group of five rats (3F) with patent catheters confirmed a significant effect of time [F (3, 14) = 11.4; P<0.001] after the start of inhalation and the post-hoc test confirmed that plasma THC was higher immediately post-inhalation compared with all subsequent time points (**Figure 10B**). Finally, analysis of plasma sampled from rats after i.p. injection confirmed a significant effect of dose [F (1, 10) = 5.92; P<0.05], but there were no significant effects of sex or the interaction of factors (**Figure 10C**).

**Figure 10:**
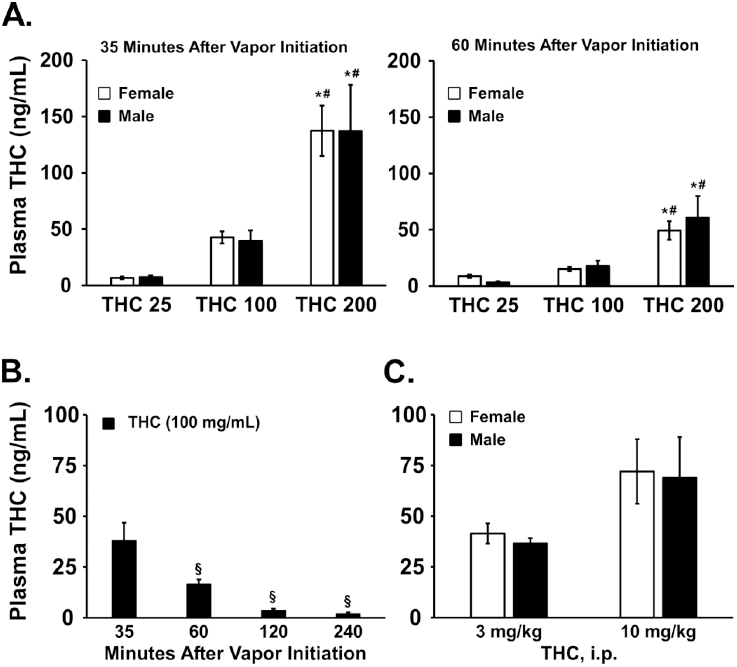
A) Mean plasma THC levels for male (N= 6 per observation; ±SEM) and female (N= 5-7 per observation; ±SEM) rats following separate sessions of vapor inhalation of the THC (25, 100 or 200 mg/mL in PG) for 30 minutes. A significant difference from the 25 mg/mL condition is indicated by * and from the 100 mg/mL condition by #. B) Mean (N= 5, 3F; ±SEM) plasma THC levels after a single session of vapor inhalation of THC (100 mg/mL) for 30 minutes. C) Mean plasma THC levels for male (N= 3 per observation; ±SEM) and female (N= 4 per observation; ±SEM) after a single injection of THC (3, 10 mg/kg, ip). A significant difference from the 35 minute time point is indicated by §.

## Discussion

This study shows that similar effects are observed in male and female rats after inhalation of Δ^9^-tetrahydrocannabinol (THC), cannabidiol (CBD) and their combination using a recently described (Nguyen et al. 2016b) e-cigarette based vapor technology. The data confirm a similar degree of hypothermia, hypoactivity and antinociception are produced in each sex after THC inhalation and none of these effects differed across estrus and diestrus phases of the estrous cycle in female rats. The data also showed that similar plasma THC levels are produced in male and female rats under identical inhalation conditions, regardless of a significant difference in body weight, consistent with the behavioral and thermoregulatory similarity across sexes. The THC inhalation conditions were chosen based on our previous study (Nguyen et al. 2016b) which showed that plasma THC levels 30 min after initiation of vapor THC (200 mg/mL) inhalation reach 176 ng/ml, and 69 ng/mL after 40 min inhalation of a crude extract estimated to contain 116 mg/mL THC in the PG. Plasma THC concentrations in human vary depending on the potency of marijuana and the manner in which the drug is smoked and studies have found peak THC plasma concentrations of 62 vs.162 ng/mL using ad libitum and paced inhalation procedures, respectively (Hartman et al. 2015; Huestis et al. 1992). The present work confirms plasma levels of about 40 ng/mL immediately after 30 minutes inhalation of 100 mg/mL THC and 137 ng/mL after inhalation of 200 mg/mL. Additional studies found a small decrease in body temperature after the inhalation of CBD alone and an interactive effect when THC and CBD were inhaled simultaneously at threshold concentrations (**Figure 6**). CBD decreased body temperature in both male and female rats and in a dose-dependent (duration of inhalation of CBD 100 mg/mL; 100 vs 400 mg/mL for 30 min) manner although the CBD effect was less incremental across dose in females and of much smaller maximum extent compared with the effects of THC in both sexes. The additive hypothermic effects of THC and CBD were observed in males and females (and hypolocomotor effects in males) when administered at a 1:4 THC:CBD ratio but no interactive effects were confirmed for a 2:1 THC:CBD ratio in male rats. Finally, there was no evidence of plasticity in the response to inhaled THC given the dosing schedule, since the magnitude of hypothermia produced by 100 mg/mL was identical from the first experiment to the final combination experiment (**Figure 8**). Thus this model produces repeatable THC exposure that is congruent with human exposure.

Although our prior study (Nguyen et al. 2016b) found a significant sex difference, with female rats apparently more sensitive to the hypothermic and hypolocomotor effects of THC inhalation, this study did not confirm a sex difference in terms of the initial magnitude of hypothermia. It may be that the former finding was related to a series of i.v. THC challenges completed in both groups prior to the vapor inhalation which led to differential plasticity of the hypothermia response, although prior work suggests female rats develop greater tolerance than males even at a lower mg/kg injected dose (Wakley et al. 2014b). Alternately it may be the case that the limitation of the prior female group to N=5 resulted in an effect of individual differences by chance. In the present study, the THC-induced hypothermia did appear to last slightly longer in the female rats (**Figures 1, 6**) which may represent increased female rat sensitivity.

The lack of a sex difference in hypothermia in these results may only appear to contrast with some prior results because of the wide dose range utilized. For example, hypothermia was greater in magnitude in female versus male rats after i.p. administration of THC at 100-176 mg/kg, but was equivalent from 1-30 mg/kg (Wiley et al. 2007). Plasma THC levels equivalent to those found in human smokers were produced by 10 mg/kg, i.p. in the present study (**Figure 10C**) suggesting anything above 30 mg/kg, i.p. in a rat may be a poor match for the human condition. Since CB1 receptor densities in the hypothalamus (associated with thermoregulation) vary across the estrous cycle in female rats (Rodriguez de Fonseca et al. 1994) it may be the case that sex differences would be obscured if estrus stage is not taken into account. However in the present study the magnitude of THC-induced hypothermia, did not vary substantially between estrus and diestrus phase (**Figures 2,3**). The possibility of faster recovery of temperature and activity during estrus observed in the higher-dose (50 mg/mL) experiment was not confirmed in the lower-dose (25 mg/mL) experiment, despite a slower initial development of the response and a quicker resolution compared with the 50 mg/mL THC inhalation.

Acute suppression of locomotor activity following either THC or CBD vapor inhalation in this study was of approximately similar magnitude across both sexes, similar to a prior report of no sex differences in the locomotor effects of 1-176 mg/kg, THC, i.p. (Wiley et al. 2007). This lack of a sex difference was, however, discordant with another study showing that 30 mg/kg THC, *s.c.*, decreased locomotor activity in male but not female rats (Marusich et al. 2014) as well as with other findings that female rodents are *more* sensitive to the acute locomotor effects of cannabinoids compared with males (Tseng and Craft 2001; Wiley 2003). These latter differences, however, occurred across all stages of the estrous cycle of the females, suggesting that hormonal levels were not the primary mediators of these differences (Tseng and Craft 2001; Tseng et al. 2004). Ovarian hormones do not modulate THC -induced locomotor suppression (Wakley et al. 2014a) and adolescents of both sexes showed comparable locomotor effects (Romero et al. 2002). In accordance with those studies, the present study showed no estrous cycle-dependent differences in THC effects on locomotion activity in females between the estrus and diestrus stages.

The present study also found that the inhalation of THC produced an anti-nociceptive effect in both female and male rats, with a sex by drug condition interaction confirming a larger effect in females (**Figure 9**). As one minor caveat, response latencies were longer in males than females and two males out of eight passed the 15 seconds cut-off time in the PG condition at 48°C; four of eight females and 5 of eight males reached the cutoff in the THC condition at the same temperature. The imposed ceiling on the maximum latency might therefore have under-represented the THC effect on males. Nevertheless, these results are reasonably consistent with previous studies in which cannabinoids were more potent in anti-nociception in female than in male rats (Wakley and Craft 2011; Wakley et al. 2014b). As with the temperature response, the anti-nociceptive effects of THC inhalation were identical across estrus and diestrus stages in this study which is similar to a prior finding after i.c.v. THC (Wakley and Craft 2011). These results were further supported by another study in which THC-induced tail withdrawal anti-nociception was not altered by the administration of estradiol, progesterone, or the combination of both hormones (Wakley et al. 2014a) and overall, THC -induced thermal anti-nociception does not appear to be sensitive to ovarian hormone modulation (Craft and Leitl 2008; Craft et al. 2012; Tseng and Craft 2001; Wakley and Craft 2011; Wakley et al. 2014a).

Cannabidiol has variously been shown to increase or oppose the behavioral effects of THC on temperature responses in rodents when administered by intraperitoneal injection, as recently reviewed (Boggs et al. 2018). Oppositional effects may depend on CBD doses eightfold higher than the THC dose (Zuardi et al. 2012) or on significant offset in the time of administration, neither of which conditions appear consistent with the smoking or vaping of cannabis or cannabis extracts. In a previous study, we found that CBD, when administered (i.p.) either simultaneously or as a pretreatment was ineffective to decrease the effects of THC on activity and thermoregulation and indeed potentiated the THC effects (Taffe et al. 2015). In a mouse model neither intravenous nor inhaled CBD reduced the effects of THC or inhaled marijuana smoke in the tetrad test, however, higher doses of CBD potentiated the anti-nociceptive effects of a low dose of THC and significantly elevated THC blood and brain levels (Varvel et al. 2006). Similarly, higher doses of CBD (10 or 50 mg/kg, i.p.) exacerbated the effects of low dose THC (1 mg/kg, i.p.) on activity, thermoregulation and spatial memory (Hayakawa et al. 2008). The present study likewise did not find oppositional effects of CBD inhalation on the thermoregulatory or locomotor effects of THC inhalation in male and female rats. There was no impact of CBD administered at a 2:1 THC:CBD ratio in male rats and an *additive* effect on hypothermia when administered at a 1:4 THC:CBD ratio in male or female rats at either threshold (**Figure 6**) or robust (**Figure 7**) THC inhalation concentrations. Interestingly, the inhalation of CBD by itself significantly reduced body temperature of both male and female rats. This contrasts with our prior finding of no thermoregulatory effect of i.p. injection of CBD in male rats (Taffe et al. 2015) and a report of no hypothermia after i.v. CBD in male mice (Varvel et al. 2006) which suggests there may be significant differences in the effects of CBD itself that depend on the route of administration. Since CBD-containing e-cigarette liquids are already appearing on the retail market (Peace et al. 2016), a rodent model is of significant use to further evaluate *in vivo* impacts of inhaled CBD in the future.

Male and female rats exhibited the same degree of hypothermia and the same plasma THC levels after vapor inhalation despite a substantial difference in body size. In particular, the comparison of the effect of identical dosing conditions (100 mg/mL THC) at two different time points across the study (**Figure 8**) shows identical magnitude of initial hypothermia across sex and across time. The females were 61% as heavy as the males at Time 1 and 52% as heavy at Time 2. Females increased by 43% from Time 1 to Time 2 whereas the males increased by 66% and the females at Time 2 were 86% as heavy as the males at Time 1. While total ventilation (Doperalski et al. 2008) and cortical blood flow (Roof and Hall 2000) of age-matched male and female rats are not different, female blood volume (Probst et al. 2006) and brain weight (Bishop and Wahlsten 1999) are lower than that of males. Therefore it might have been predicted that females would have experienced a higher effective dose. As reviewed above, where sex differences have been reported following parenteral injection on a mg/kg weight adjusted basis, female rats tend to be more sensitive than male rats. In this study, no major sex differences were observed across dosing conditions that produced significant dose-dependent effects, nor did they differ in plasma THC concentration. Thus it is most parsimonious to conclude that any sex-dependent differences in effective brain concentrations that may have been reached at a given dosing condition were not large enough to compromise the overall conclusion of minimal sex differences in response to inhaled THC, *in vivo*.

In conclusion, we have shown that the sex of the rat does not modify the general pattern of hypothermic or hypolocomotive effects of inhaled THC, CBD or their co-administration, nor the anti-nociceptive effects of THC inhalation. The analysis of plasma THC further confirmed similar exposure to THC across sexes following vapor inhalation under similar conditions. In addition, there was no difference in the female rats’ response to the inhalation of THC between estrus and diestrus stages across thermoregulatory, activity and nociceptive measures. These results contrast with some prior results reported for parenteral injection of THC or other cannabinoids. It may be the case that this new model of e-cigarette type vapor inhalation provides an improved technique for the evaluation of cannabinoid effects in rodent pre-clinical models, possibly because the *in vivo* effects have relatively rapid onset and offset relative to injected doses that produce similar magnitudes of effect (Nguyen et al. 2016b; Taffe et al. 2015).

## Acknowledgements

Funding and Disclosure: This work was funded by support from the United States Public Health Service National institutes of Health (R01 DA024105, R01 DA035482 and R44 DA041967) which had no direct input on the design, conduct, analysis or publication of the findings. MJP was supported in part by T32 AA007456. MC is proprietor of La Jolla Alcohol Research, Inc and PI of the SBIR grant (R44 DA041967) supporting further commercialization of the inhalation equipment. The authors declare no additional competing financial interests for this work. The authors are grateful to Mr. Howard Britton for prototyping and fabrication of inhalation equipment and to Arnold Gutierrez, Ph.D. for comments on the manuscript draft. This is manuscript #29416 from The Scripps Research Institute.

